# What goes up must come down Biomechanical impact analysis of jumping locusts

**DOI:** 10.1101/225672

**Authors:** Simon V. Reichel, Susanna Labisch, Jan-Henning Dirks

## Abstract

Many insects are able to precisely control their jumping movements. Previous studies have shown that many falling insects have some degree of control of their landing-orientation, indicating a possible significant biomechanical role of the exoskeleton in air righting mechanisms. Once in the air, the properties of the actual landing site are almost impossible to predict. Falling insects thus have to cope mostly with the situation at impact. What exactly happens at the impact? Do locusts actively ‘prepare for landing’ while falling, or do they just ‘crash’ into the substrate?

Detailed impact analyses of free falling Schistocerca gregaria locusts show that most insects typically crashed onto the substrate. There was no notable impact-reducing behaviour (protrusion of legs, etc.). Independent of dropping angle, both warm and cooled locusts mostly fell onto head and thorax first. Our results also show that alive warm locusts fell significantly faster than inactive or dead locusts. This indicates a possible tradeoff between active control vs. reduced speed. Looking at the morphology of the head-thorax connection in locusts, we propose that the anterior margin of the pronotum might function as a ‘toby collar’ structure, reducing the risk of impact damage to the neck joint. Interestingly, at impact alive insects also tended to perform a bending movement of the body.

This biomechanical adaptation might reduce the rebound and shorten the time to recover. The adhesive pads also play an important role to reduce the time to recover by anchoring the insect to the substrate.

## I. Introduction

For many insects both flight and jumping movements are means of efficient and fast locomotion. In general, flight and jumping movements can be subdivided into three major phases: take-off, aerial phase and landing. However, whilst landing after flight is mostly a predictable movement which is actively controlled by the insect (Borst, 1986; Manzanera and Smith, 2015), landing after a jumping movement has additional challenges. Although the initial trajectory of the takeoff and even the orientation of the body might be controlled by several jumping insect species (Alexander, 1995; Faisal and Matheson, 2001; Heitler, 1974; Rothschild et al., 1972; Sutton and Burrows, 2008, 2011), external factors such as gusts of wind or unforeseen obstacles make it extremely difficult to reliably predict the mechanical properties of the landing site (Bennet-Clark and Alder, 1979). An insect landing after a jump thus must be able to cope with a large variety of possible surface properties. Surfaces could be elastic or inelastic, smooth or covered in obstacles, plain or tilted, and each possible combination thereof. As reliable information about the physical properties of the impact side can not necessarily be acquired from visual cues during the limited timeframe of the late aerial phase, falling insects have to cope with the situation at impact.

For many jumping insects such as Cercopoidea (froghoppers) or Siphonaptera (fleas) with a very small body mass, the impact on the substrate is probably not an important factor to take into account in respect to damage to the exoskeleton. However, for jumping insects with a body mass several orders of magnitude larger, such as locusts, it is likely to be more important to land in a controlled way to reduce the risk of damage of the exoskeleton. In addition, a successful and controlled landing movement resulting in a quick upright and stable body posture minimizes the time to prepare for the next jump. This would increases the chance of successful escape of a large and traceable insect from a predator.

The air-righting mechanisms during flight and falling are relatively well understood (Robert and Rowell, 1992; Weis-Fogh and Jensen, 1956; Zarnack, 1978). A previous study by Faisal and Matheson, (2001) has shown that locusts use combinations of active and passive movements of their legs and wings to turn themselves mid-air, which significantly increases the chances to land upright, even when dropped upside down (Faisal and Matheson, 2001). Locusts are also able to control tumbling during the aerial phase (Cofer et al., 2010).

However, despite increasing knowledge about the aerial phase, the actual biomechanics of the impact itself are still unclear. Do locusts just crash into the substrate and rely on their stable exoskeleton to deal with the impact forces? If initial ground contact is not random, some sort of landing control or energy absorbance behavior might be present to ensure a quick recovery and stable position independent of the properties of the substrate. Active control of landing could involve similar actions as described for air righting such as usage of legs or wings. If there are passive control effects involved, the exoskeleton of the locust might provide interesting features that favor landing in the upright position, such as aerodynamic features or the performance of the adhesive pads.

To examine active and passive control features, locusts were tested in three different living stages on a variety of landing parameters to reveal differences in landing performance. The information acquired was then used as an biomimetic inspiration for possible applications in the technical context.

## II. Materials and Methods

### 1. Insect specimen

Adult male and female *Schistocerca gregaria* locusts were kept at room temperature (12 h day at 24 °C, 12 h night at 19 °C) and an relative air humidity of 50 – 60 %. The insects were fed fresh food *ad libitum*. Locusts with missing legs or defect wings were excluded from the experiments. For automated motion tracking the center point between eyes and mandibles on the frons and the central tip of the abdomen were chosen as markers. Each group included twelve locusts equally distributed between male and female locusts.

### 2. High-speed recordings

A high-speed camera (*Fastcam APX RS MONO* by Photron, Tokyo, Japan) with a 50 mm objective was used to record the landing performance at 1500 fps and bright ambient light conditions (about 20.000 lx). The camera was aligned to record front and side view of the falling locust as well as the ground (smooth glass) using a surface mirror at 45°. To correct for lens distortions at different positions within the video frame, the setup was calibrated using a custom-made script utilizing the *Python* bindings for the calibration algorithms of *openCV* (Bradski, 2000). The highest reprojection error recorded was 0.13.

A metal stand was used to ensure a constant dropping height of 0.6 m and a fixed dropping angle. To prevent any effect of pressure induced thanatosis (Faisal and Matheson, 2001), the insects were carefully held by grabbing the folded wings with thumb and index finger right behind their base. All insects were dropped without allowing the locust to push against any object and thereby adding unwanted torque to its fall.

The first set of locusts was dropped parallel to the ground, the second one in a positive 45° pitch angle (abdomen upwards) and the third in negative 45° direction (abdomen downwards). Based on the previous study by Faisal and Matheson, a dropping height of 0.6 m was selected. At this height, free falling *S. gregaria* locusts are able to land upright (Faisal and Matheson, 2001). The high-speed recordings of the locust landing were performed at 19 °C with 50 — 60 % relative air humidity. For each set every locust was tested alive and warm (body temperature approx. 24 °C), alive yet cooled (approx. 4 °C) and finally dead. Usually, dead insects desiccate rapidly, which affects the biomechanical properties of the cuticle (Dirks and Taylor, 2012). Previous studies however have shown that freezing insect cuticle neither significantly affects its static nor dynamic biomechanical properties (Aberle, Jemmali and Dirks, 2017). Hence locusts were sacrificed by freezing them to —20 °C, then thawing the dead insects to room temperature immediately before the experiments. The experiment was conducted on one locust at a time to minimize the cooling-down and warming-up of the insect respectively. None of the experiments took longer than 15 minutes for warm locusts and not longer than 3 minutes for cool locusts.

The end of the landing phase was defined as the moment when a locust reached a stable stance on the substrate, with the femora of the hind legs orientated at approximately 40 degree from the ground and the tibiae fully flexed (Faisal and Matheson, 2001) or no further body movement was visible. The timeframe between impact and reaching a final stable body posture was defined as the *landing duration*. For dead locusts the end of landing phase was the moment when it ceased bouncing and gliding on the ground surface.

To investigate the effect of wings on impact speed and angles, paired experiments were performed on locusts with and without front and hind wings. Locusts were also dropped parallel to the ground undamaged first (warm), after removal of the wings (warm) and finally in cooled body stage. The effect of adhesive pads was tested by removing all tarsal segments from front, middle and hind legs.

### 3. Tracking and reconstruction

The calibrated high-speed recordings were automatically analysed using pattern motion tracking scripts (Blender, version 2.78a, Blender Online Community, 2016). Markers were tracked in front and side view as provided by the mirror. Manual interpolation or offset tracking of the marker position was required when the patterns were obscured by wings, legs or rotation of the locust. The marker positions were manually checked and adjusted forth and back in time.

Four main parameters of the fall were reconstructed from the video data: impact angle, impact speed, landing duration and body part of first contact.

A three-dimensional vector between head and abdomen markers of the locust was reconstructed using the *x* and *z* coordinates from the front and the *y* coordinates from the side view. This vector was used to calculate the locust angle relative to the planar ground surface assuming nearly orthogonal coordinate space from the image data. Angles generally were defined between the ground plane (x-y plane) and this longitudinal body vector. The impact angle was averaged over the last three frames before impact. Consequently, a positive impact angle means the locust hit ground head first opposed to abdomen first for a negative impact angle.

The same coordinates were used to reconstruct the speed of the locust. The impact speed was calculated from the movement of the head marker during the last three video frames prior to impact moment in all three dimensions. An order 3 Butterworth low-pass filter (0.15 Hz) with initiation on the mean of the first 10 speed values was applied to reduce noise.

Assuming the kinematic equations of speed and acceleration the theoretical impact speed of a locust, ignoring any aerodynamic effects, was calculated as

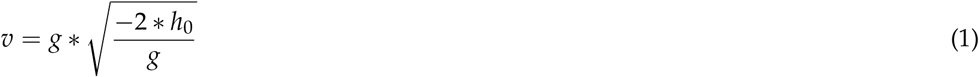

with the gravitational constant g = –9.81 m/s^2^ and h_0_ = 60 cm as the dropping height. This simplification results in a maximum velocity of about 3.43 m/s of a falling locust at impact.

The *first contact* described the part of the locust body which first touched the ground. In this respect both antennae and head of the locust were considered a *head* contact, whilst *full body* describes an impact with an indistinguishable part of the thorax or abdomen. Such first contacts only occurred when locusts were aligned horizontally before impact.

Additional qualitative observations of the falling behavior were also recorded. In many cases characteristic bending movements of the abdomen were observed. The abdomen bent away from the longitudinal body axis after impact (see figure 6). When this movement exceeded the threshold of 20° from the body’s main axis, the abdomen was considered as *bent*.

### 4. Statistics

Statistics were performed using *R* (version 3.3.2, R Core Team, 2016). The index of the test statistics always represents the degrees of freedom (numerator, denominator if applicable). Shapiro-Wilk tests were used to test for normal distribution. As two-sample test for homoscedasticity the F test was used. Either Student’s t-test or Welch’s t-test were chosen according to the prerequisite of variance. For non-normally distributed data the Wilcoxon rank sum test was used.

To compare multiple samples different tests were chosen, depending on the prerequisites. For a one-way ANOVA both normality and homoscedasticity were met. For the Kruskall-Wallis test only homoscedasticity was required (in short Kr-Wa). For the Welch-corrected ANOVA only normality was met (in short Welch’s ANOVA). Multi-sample comparison of variance was performed with Bartlett’s test for normal-distributed samples and Levene’s test for nonnormal data. When significance occurred Dunn’s test was used to distinguish the different subgroups for ANOVA and Kruskall-Wallis tests. For the Welch-corrected ANOVA a pairwise t-test with Holm-Bonferroni correction was used to distinguish the subgroups (in short Holm t-test).

A significance level of *α* = 5% was used in all tests. If not stated otherwise the provided values show the median with the median absolute deviation (*median* ± *mad*). If mean values are used the standard deviation is given instead (*mean* ± *sd*). Boxplots show the median and the last datapoint within 1.5 boxlength as whiskers. Outliers were identified using the Tukey’s range test and removed from the analysis for each test set individually. Additionally, for pairwise t-tests the measurements referring to an outlier were removed from the paired groups, too.

## III. Results

### 1. Pre-impact

#### Impact speed

A typical head speed progression of a living, warm locust over a whole video sequence is shown in figure 1. Given a dropping height of 0.6 m, the still accelerating falling locusts reached their highest speed just before impact. After impact, the relative speed was ‘positive’ as most of the insects showed rebound movements from ground. As the locusts reached their stable end positions the head speeds leveled out at zero.

**Figure 1:**
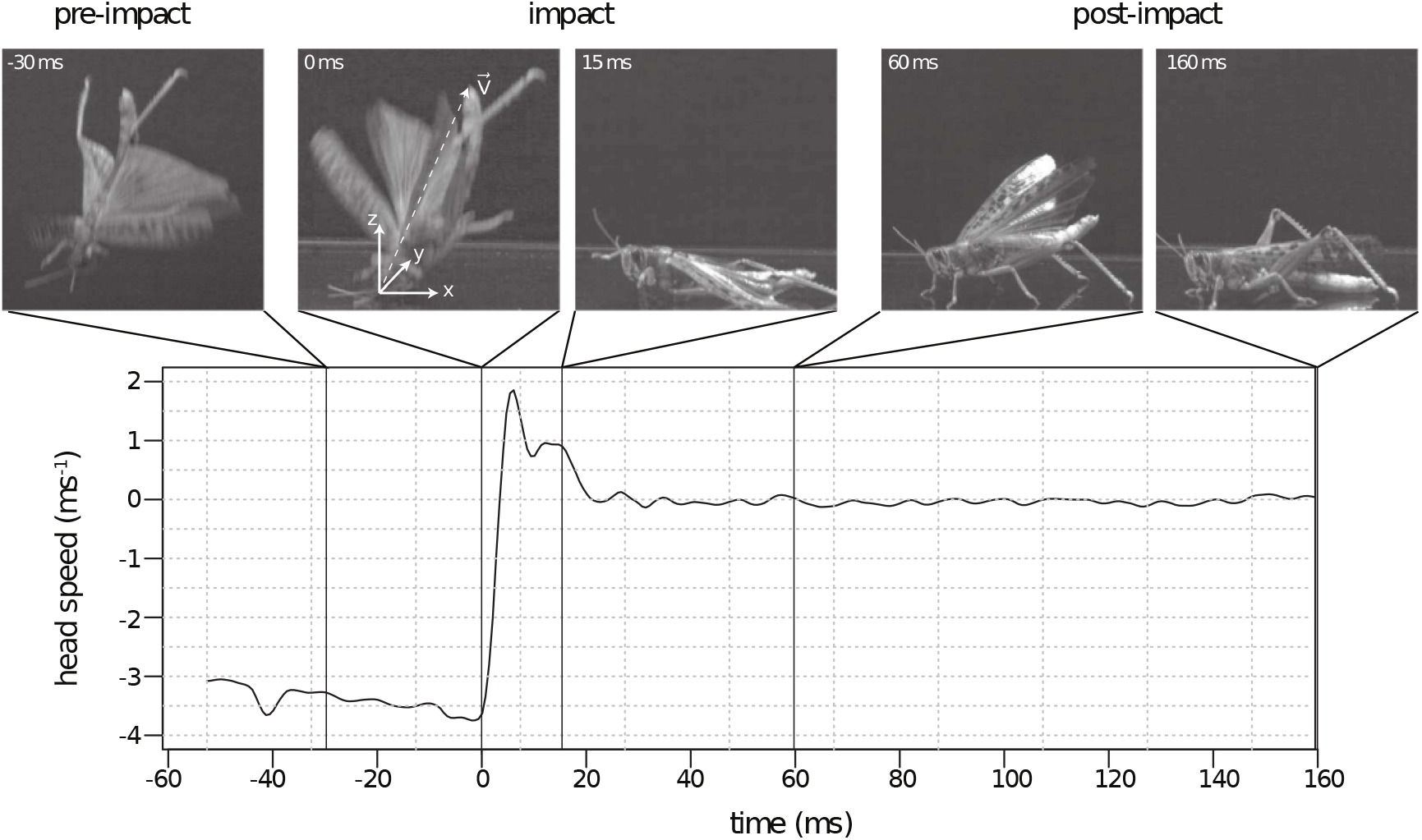
Speed of the locust head over time for a typical recording of a living, warm locust falling from a height of 0.6 m onto a planar glass substrate. The frames show the locust at characteristic stages of the landing movement: pre-impact (ca. –30 ms), impact (0 to 30 ms) and post-impact (> 30 ms). During the late pre-impact phase often the front and hind wings were actively moving. Warm locusts tended to crash head first onto the substrate. This was followed by a characteristic bending movement of the abdomen (see suppl. video). A stable final stance was typically reached after about 150 ms.

Our results show a significant effect of activity level on impact speed (see figure 2 A). Warm insects at planar drop angle had a significantly higher impact speed (3.73 ± 0.13 m/s) than dead insects (3.33 ± 0.07 m/s) (Kr-Wa H_2_ = 14.701, *p* < 0.001 with Dunn *p* < 0.05). In comparison, the impact speed of cold insects (3.44 ± 0.07 m/s) was still significantly larger than the impact speed of dead insects (Dunn *p* < 0.05).

**Figure 2:**
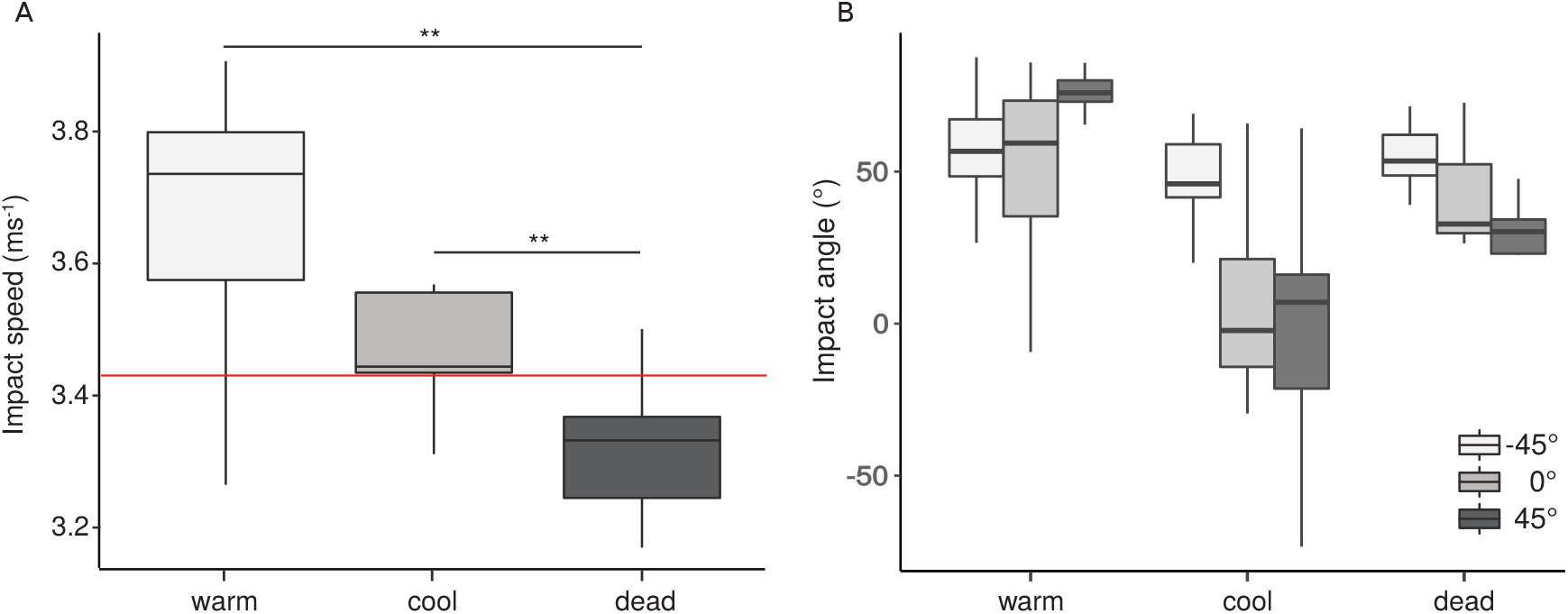
A) Impact speed for three different states of activity. Both the warm and the cooled locusts fell significantly faster than the dead locusts. The theoretical maximum impact speed of 3.43 m/s from a height of 0.6 m is marked in red. Interestingly, the warm insects fell faster than the expected maximum falling velocity. This is likely a result of acceleration by flapping the hind wings during the final stages of the landing (see 1 first frame). B) Impact angle over locust state of the three different drop angles. Warm locusts were able to correct for different dropping angles and rotate their bodies to ensure a head-impact. Cooled insects showed a notably higher variability of impact angles in comparison to warm insects. Planar and abdomen-first drops resulted in less steeper impacts. Dead locusts again showed a much smaller variability of impact angles. Even at positive, abdomen-first dropping angles, the bodies of dead locusts rotated in the air, resulting in a head-first landing.

Changing the drop angle either ±45° from the horizontal plane in general reduced the impact speed. Warm insects with negative drop angle (abdomen first) had a significantly lower impact speed (3.28 ± 0.21 m/s) than the positive or planar drops (Kr-Wa H_2_ = 16.848, *p* < 0.001 with both Dunn *p* < 0.001).

#### Impact angle

The effect of dropping angle on impact angle for all activity states is shown in figure 2 B. At planar drop angles, the activity state significantly affected the impact angles (Welch’s ANOVA *F*_2,17.69_ = 75.032, *p* < 0.001 with all Holm t-test *p* < 0.05). The steepest mean impact was found for warm locusts with 75.43 ± 7.80°. Cold insects instead had a much lower mean impact angle of 2.32 ± 38.69°. Interestingly, the results of dead animals lay between those two, with a mean impact angle of 31.72 ± 8.67°.

At positive drop angle (head first) the cool locust stance showed (—2.27 ± 25.03) significantly differences to both warm and dead stance (Kr-Wa H_2_ = 11.559, *p* < 0.01, with both Dunn *p* < 0.01). Warm (59.37 ± 22.82°) and dead (32.77 ± 7.60) did not differ significantly (Dunn *p* = 0.3862). However, at —45° dropping angles the locust activity state did not significantly affect impact angles (ANOVA F_2_ = 2.235, *p* > 0.05).

#### Effect of wings

The effect of wings on impact speed and angles was analysed using paired experiments on locusts with and without front and hind wings. Our results show that the impact speed of a locust with front and hind wings was not significantly different to the same locust without wings (Paired *t*-test *t*_9_ = –0.1002, *p* = 0.92, see figure 3 A) when dropped parallel to ground. However, the impact angle did change significantly when the wings were removed (Paired *t*-test *t*_9_ = –2.4853, *p* < 0.05). Locusts without wings landed almost parallel at a mean angle of –3.44 ± 51.09° while winged locusts had a head-first impact with 39.28 ± 30.89° (see figure 3 B).

**Figure 3:**
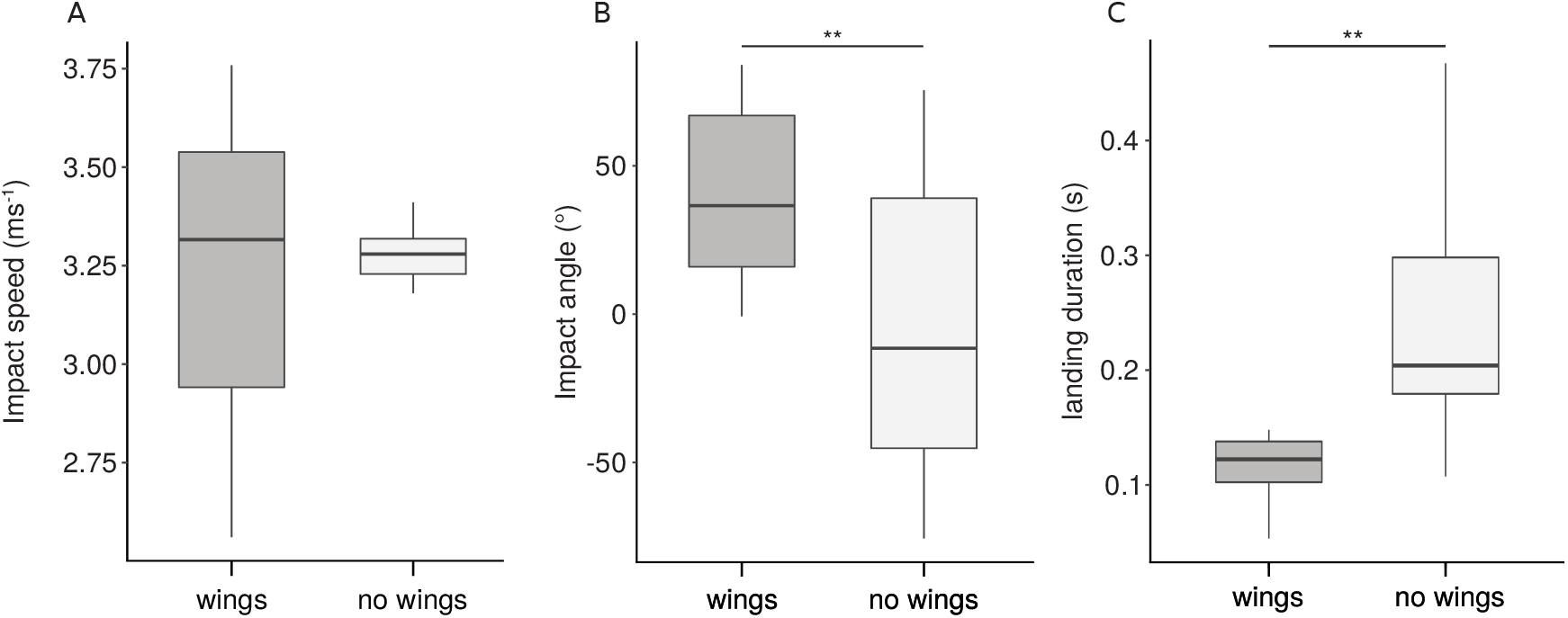
A) Head speed of warm locusts with and without wings before impact after a horizontal drop. There was no significant difference in impact speed between locusts with and without wings. B) Body angle of warm locusts with and without wings before impact after a horizontal drop. Locusts with wings had a significantly steeper impact angle, landing head first. C) Landing duration of warm locusts with and without wings. Locusts with wings had a significantly shorter landing duration, thus would be able to perform a consecutive escape jump within a shorter period of time.

### 2. Post-impact factors

#### First contact

The relative distributions of the body parts which touched ground first are shown in figure 4. Pooling the different activity states, at plane drop angle 58 % of the locusts hit the ground head first. Front and middle leg were involved at almost equal proportions of 14 % and 17 % respectively. The abdomen (6 %), full body (3 %) and hind leg (3 %) occurred rarest. At negative drop angles head and front leg represented most of the contact points with 64 % and 31 % respectively. At positive drop angles the head proportion was slightly lower (42 %) than the planar drops, and front or middle leg occurrence was increased to 22 %, respectively. The distinguishable body states can be seen in more detail in figure 4.

**Figure 4:**
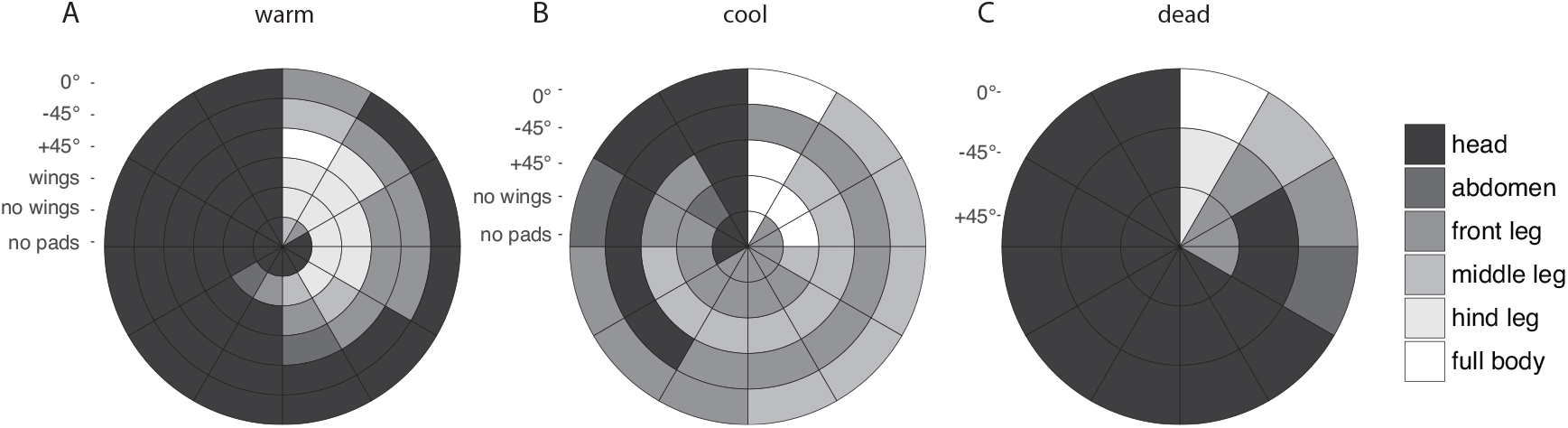
Body-parts of locusts with first contact at impact to the substrate under different prerequisities. While warm locusts tend to land head first, cooled insects showed a notably higher variety of first contacts. Dead locusts, however, showed a tendency to land head first. These results correlate well with the effect found in the impact angles and indicates the presence a passive descend-control mechanism, which in cooled, yet alive insects might be affected by uncontrolled or delayed body movements.

Combining the first contact data with impact speed, impact angle or landing duration did not provide enough data to be statistically solid, however, the general tendency indicates, that the contact point changes with the impact angle. As figure 4 illustrates, cool insects showed a notably higher variability of first-contact-points than the warm insects. The head was less involved, while front and middle legs touched the substrate much more often. Interestingly, the dead insects again showed a more stereotypic pattern similar to the warm locusts, as the headwas the primary point of first-contact at impact. Legs or abdomen were rarely involved.

#### Landing duration

The effect of activity state and dropping angle on landing durations of the locusts is shown in figure 5 A. The dropping angle had no significant effect on the landing duration, the activity state however significantly affected the time to recover. For planar drop angle the warm locusts had a significantly lower mean duration of the landing (0.136 ± 0.023 s) than the cool (0.276 ± 0.080 s) or dead (0.249 ± 0.061 s) insects (Welch’s ANOVA *F*_2,15.73_ = 28.089, *p* < 0.001 with paired Holm *t*-test *p* < 0.001). The same tendency was visible at positive and negative drop angle. Between the durations of warm locusts was no significant difference for the three dropping angles (Welch’s ANOVA with all Holm t-test *p* > 0.05). Same applies to the cold (Welch’s ANOVA *F*_2,19.37_ = 0.171, *p* = 0.844) and dead locusts (ANOVA *F*_2_ = 0.1, *p* = 0.905).

**Figure 5:**
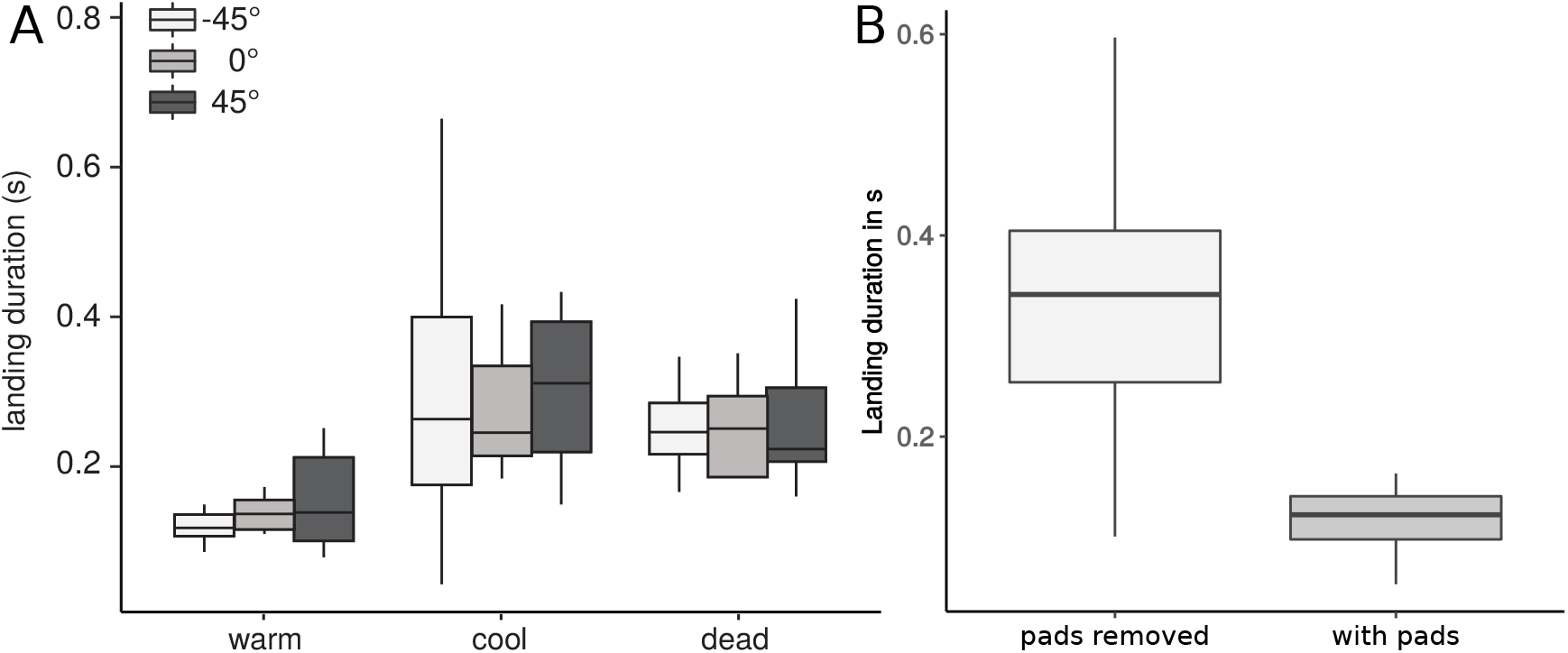
A) Landing duration of warm locusts at different activity states and dropping angles. Warm locust took significantly less time to reach a final resting position than cool or dead insects. The dropping angle however had no significant effect on the duration of the landing, irrespective of activity state. B) Landing duration for warm locusts with and without adhesive pads. Locusts with their tibiae removed had a significantly longer landing duration.

In contrast to the warm and cooled locusts, which all sooner or later reached a stable upright resting position, dead locusts instead showed a variety of different final states after the mostly head-first impacts. At planar drop angle 92 % rested sideways and only 8 % upright. When released with a positive drop angle 84 % rested sideways, 8 % backwards or upright each. However, at negative drop angle only 59 % rested sideways, while 33 % rested upright (8 % backwards).

Warm locusts, which had their tibiae removed, took significantly longer to reach a stable state after impact from planar drop angle (Welch’s t-test *t*_10.876_ = 5.112, *p* < 0.001). The mean landing duration of insects without adhesive pads was 0.335 ± 0.137 s compared to 0.118 ± 0.030 s for locusts with adhesive pads (see figure 5 B).

#### Dissipation of kinetic energy

Detailed analysis of the high-speed recordings after impact revealed interesting characteristic biomechanical behavior of the abdomen. After impact, the abdomen either stayed almost completely in its initial ‘straight’ shape or the abdomen bent upwards (‘bending’, see figure 6).

**Figure 6:**
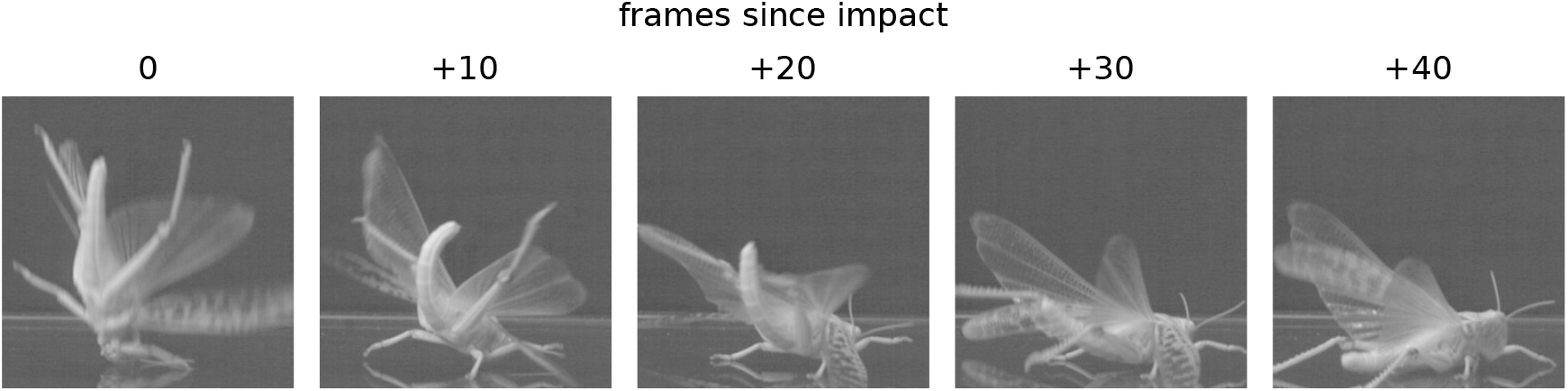
Bending sequence of a locust (positive drop angle). The delay between the shown images was ten frames each. Right after impact the abdomen bent upwards and landed with a rolling movement thereafter.

Our results show that the impact speed at planar drop angle was not significantly different between locusts that bent or kept a straight abdomen (Student’s t-test *t*_34_ = –1.642, *p* > 0.05). However, as the bending behaviour was not distributed equally between body states, these results have to be used with care. Warm locusts at planar drop angle were much more likely to show bending behavior (83/17 %) than cool (25/75 %) or dead locusts (33/67 %, see figure 7 A)). Taking all three locust states at planar drop angle into account, those showing no bending behaviour after impact took significanlty longer to recover (Welch’s t-test *t*_25_._26_ = –5.0745, *p* < 0.001). However, the same rule as for the dropping angles applies here, and the results need to be seen in context of the body states and the involved distributions. The landing duration can be seen in figure 7 B).

**Figure 7:**
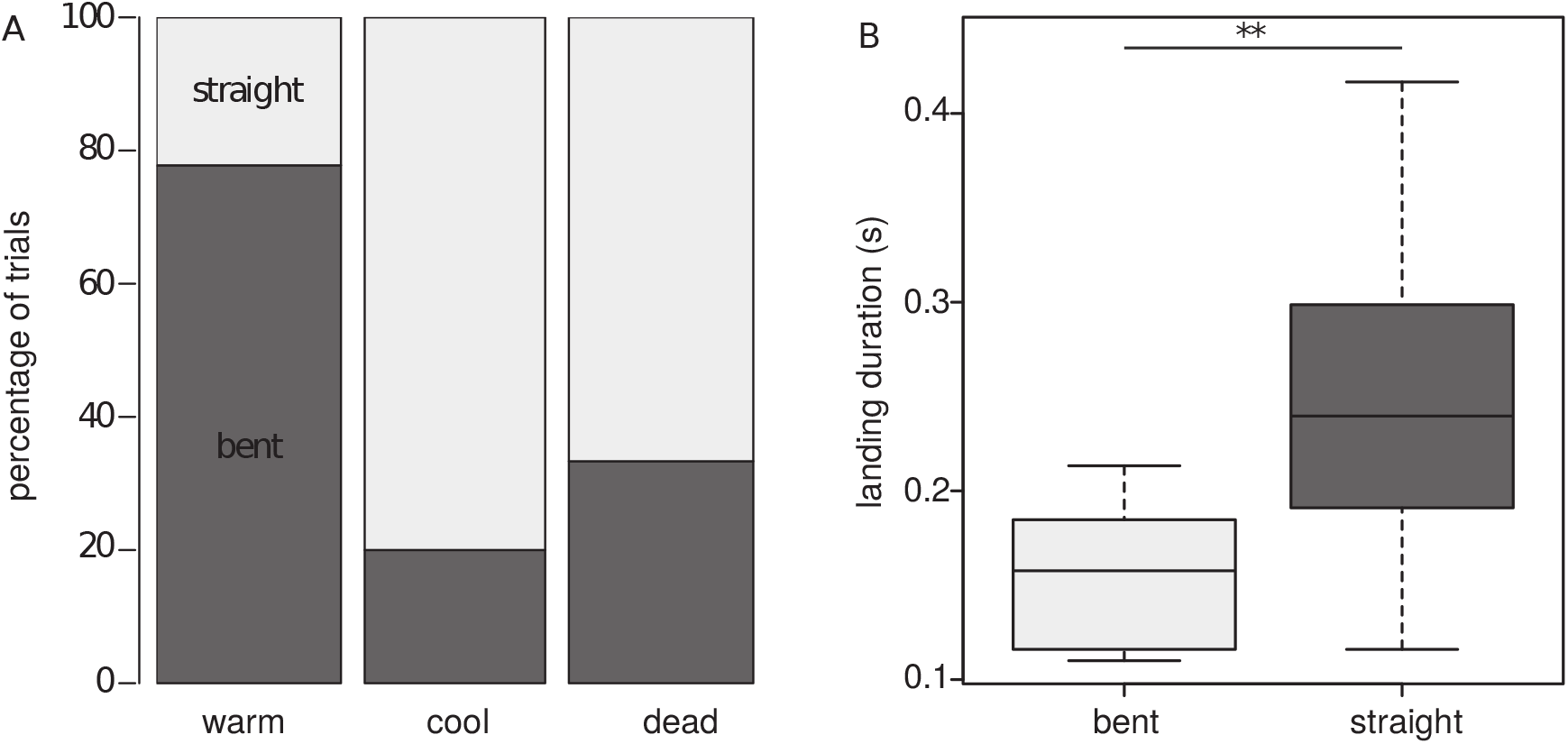
A) Percentage of locusts showing bending movements of the abdomen at the three different activity levels (planar drop angle). Warm locusts show a notably higher degree of bending than cooled or even dead locusts. Interestingly, also dead locusts showed bending of the abdomen to some degree, indicating a passive component of this behaviour, which might be enhanced by activity of the locusts. B) Landing duration of locusts at planar drop angle which showed bending of the abdomen. Locusts with bending showed a significantly shorter landing duration than locusts with no bending movement of the abdomen.

## IV. Discussion

The biomechanics of the takeoff before flight and jump have been well studied across a variety of different insect species (Manzanera and Smith, 2015). Various sophisticated mechanisms and structural adaptations have been shown to allow for example efficient energy storage, highly synchronized leg movement and detachment from the surface (Christian, 1978; Rothschild et al., 1972; Sutton and Burrows, 2011). Locusts for example, a typical model organism for jumping biomechanics, are able to control the initial trajectory of their jump in six degrees of freedom (Eriksson, 1980; Gvirsman, Kosa and Ayali, 2016; Sutton and Burrows, 2008). During the aerial phase several insects show active and passive righting mechanisms, ensuring a controlled spacial orientation of the body (Chahl, Srinivasan and Zhang, 2016; Goodman, 1960).

Controlled and predictable landing after flight, mostly studied for various species of Calliphorinae, typically involves optical flow to estimate the distance ground, followed by preparatory movements of the legs, where the prothoracic legs are extended and slightly lifted to make first contact with the ground, whilst at the same time the meso- and metathoracic legs are lowered (Borst, 1986; Goodman, 1960). The focus of this study however was ‘unpredictable’ landing of jumping insects. Especially for insects, neither the duration of the aerial phase, nor the quality of the landing cite can be anticipated.

In insects, body temperature is obviously closely correlated with general activity and in particular flight (Krogh and Weis-Fogh, 1951; Mellanby, 1939; Taylor, 1963). The lower the body temperature of the locusts in our experiments, the less likely were any effects of active movements or control of the impact. Dead, yet not-dehydrated insects obviously had no control of the falling movement, whilst still having the same passive biomechanical properties of their exoskeleton (Aberle, Jemmali and Dirks, 2017).

### 1. Active and passive pre-impact control

Our experiments with free falling locusts at different ‘activity levels’ show that irrespective of the initial dropping angle, free falling warm locusts, with the highest degree of control, showed a characteristic impact angle of about –75°. Warm locusts mostly fell ‘head first’ onto the substrate, slightly spreading their wings and legs (see figure 2 B). These results are very consistent with results previously published by Faisal and Matheson, (2001). Dead locusts however also showed a characteristic ‘head first’ impact angle very similar to the impact angle of warm locusts, again irrespective of dropping angle. These results also directly confirm the passive aerodynamic air-righting effect of the locust exoskeleton, earlier suggested by Faisal and Matheson, (2001).

Surprisingly, the results of this study show that cold locusts fell with not only a less steeper impact angle in comparison to the active locusts, however also showed a notably higher degree of variation in comparison to both the warm and dead locusts. Hence, regarding impact angles, being dead is better than being mostly inactive. This seems surprising, however might be explained by probably uncoordinated corrective active movements of the cold locusts. Locusts extending their hind tibiae and spreading the wings were notably more likely to land upright than locusts which did not show this active behavior (Faisal and Matheson, 2001). When falling from an orientation close to the preferred impact angle (–45 vs. –75°), the locusts performed only little corrective movements. However, when falling from either 0 or +45°, the falling locusts most likely performed movements of probably legs and wings, which resulted in no or only chaotic correction of the impact angle.

Our results also show that warm locusts fell significantly faster than dead locusts (see figure 2 A) and faster than the cool ones by trend. Interestingly, our results also show that warm locusts also fell faster than predicted by a simple free-falling approximation (ignoring air resistance). This rather unintuitive observation could only be a result of active movements and indicates a possible tradeoff between active control vs. reduced speed.

As the wings presumably play an important role in the air-righting behaviour (Faisal and Matheson, 2001), it thus seems likely that rather than to decelerate their falling speed, locusts might have used their wings to gain control and at the same time speed (similar to a go-around of a landing aircraft). Indeed, locusts with wings were able to control their impact angle during free fall, whilst wingless locusts landed at almost the same impact angle they had been dropped with (see figure 3 B). This effect might be easily explained with active steering movements of the wings.

However, our experiments also show, that surprisingly the presence of wings had no significant effect on the locust impact speed (see figure 3 A). This leaves the question: where do the differences in speed based on ‘activity levels’ come from? As the simple free-fall approximation did not consider air friction or orientation, the body posture itself might have an effect on the impact speed. Another possible factor determining falling speed could be the orientation (and thus air resistance) of the legs, which has also been indicated by the studies of Faisal and Matheson, (2001). Our observations show that typically dead locusts had their legs close to the body, while warm and cool locusts often had their hind legs stretched out. Further high-speed recordings of the free fall might help to answer these open points in the future.

In summary, locusts use both active and passive mechanisms to control both impact speed and impact angle during free falling. Whilst the presence of the wings does not significantly affect the impact speed, wings do significantly affect the control of the impact angle. From the tested dropping height free falling locusts did not decelerate, which would have reduced the mechanical impact on the exoskeleton. Instead the insects rather even accelerated to gain control of the impact angle. Impact speed thus seems not to be a limiting factor to protect the exoskeleton, whilst impact angle, and thus a control of the body part with first contact to the substrate, might be. At higher dropping heights, or landing after flight, the wings might obviously play a different role in deceleration.

### 2. Post-impact biomechanics

Right after impact, locusts need to cope with the kinetic energy of the impact to avoid damage of the exoskeleton, as well as prevent falling over. One possibility of controlled active deceleration after a jump is the use of legs, which can be found in a great variety in many different animals (Alexander, 1995). However, our results show that this principle holds not true for locusts.

#### First contact

The previous experiments already showed that at impact warm locusts tend to have a tilted head-first body posture (see figure 2). The detailed post-impact analysis of the actual first body part in contact with the substrate shows that, instead of using for example the legs to absorb impact energy, the major fraction of active locusts landed literally ‘head-first’, making mostly first contact with the mouthpart region of the head (see figure 4 A). The first contact occurs rarely with any parts of the legs, thorax or the abdomen. It is also, most likely as a direct consequence of the impact angle, independent of dropping angle for warm insects. Neither the presence or absence of wings or adhesive pads did notably affect the first point of contact.

Interestingly, this effect was also found for dead locusts (see figure 4 C). This shows that landing on the head is a purely passive result of the impact angle and plays an important role in locust landing.

For cooled, and thus only partially active locusts, the analysis of first contact shows a pattern obviously correlated to the impact angle of the same insects. The locusts with different impact angles show a large variety of first contact points (see figure 4 B), with legs and abdomen involved.

One qualitative observation found in our video recordings was a characteristic small movement of the locust head in relation to the pronotum during impact. Detailed analysis of the head showed that after impact the head slightly moved backwards into the pronotum. This movement appeared shortly before any other reaction (such as bending, see below) was observed. Looking at the morphology of the head-thorax connection in locusts, it seems likely that the anterior margin of the pronotum in combination with the posterior part of the head (occiput) might function as a ‘toby collar’ structure. This structure would act as a temporary head-arresting mechanism, reducing the risk of impact damage to the relatively small neck joint.

#### Landing duration

A short landing duration, i.e. the time between impact and readiness for the next jump, plays an important role for any escaping jumping insect. The faster an escaping locust is able to perform a consecutive escape jump, the more likely it is to escape a predator.

Minimising landing duration can be achieved by combinations of two general strategies. In the first place, an insect falling on its side or back after an escape jump could try to use an efficient and fast uprighting mechanism. This mechanism has been described for locusts in detail by Faisal and Matheson, (2001) and mostly involves movements of the legs pressing agains the ground, as well as rolling movements of the body. Typically, a locust turned upside down takes 600 ms to complete this movement and become ready for a consecutive jump. A timeframe of 600 ms however is a considerable long period of time when escaping for your life. A second additional strategy would therefore be to initially land in a correct (or at least close to correct) position, which requires only little correction. In our experiments, sidewards landing only ever occurred in few trials of dead insects.

Our results show that landing duration was independent of the dropping angle (see figure 5), which is trivial as the dropping angles did not significantly affect the impact angles. Hence, all warm insects had almost the same ‘starting position’ after impact.

The typical landing duration of a free falling warm locust was about 137 ms. This corresponds very well with the typical time frame reported by Faisal and Matheson, (2001) for biomechanical preparation of the next jump after landing. The landing duration for cooled insects was obviously significantly longer (230 ms). However, for cooled insects, the landing duration also showed a notable larger degree of variability, which could be a result of less coordinated body movements.

#### Dissipation of energy

To gain control of their landing movements, falling insects need to dissipate the kinetic energy of their free fall through their exoskeleton. Otherwise, the locusts would bounce of the substrate.

The detailed analysis of the landing behavior in our experiments shows that locusts did not use any noticeable protrusive movements of their legs to dampen their impact. Such behavior could have indicated the use for example friction in their joints or dynamic damping properties of the leg muscles.

A simple and easy way to dissipate the energy and prevent any ‘rebound’ is to use an anchoring mechanism to the substrate. The adhesive pads of insects are an obvious structure to provide such a mechanism (Dirks and Federle, 2011). Typically, secure and firm attachment to smooth or rough substrates does not require any active controlled movements of the insect foot (Endlein and Federle, 2008), hence adhesive pads could passively operate in ‘real time’ immediately after impact. Indeed, our results show that the landing duration increased when the adhesive pads were removed from the legs. The adhesive pads, and their ability to attach to a substrate, thus play an important role in determining the landing biomechanics of jumping insects.

The observed difference in landing duration between the warm and cooled locusts however also shows that the adhesive pads (which were present in both groups) are not the only factor to control the dissipation of kinetic energy. Instead, our results indicate the presence of a dissipation mechanism more closely linked to the body temperature and thus activity of the insect.

Indeed, our results show that just after impact, alive insects tend to perform a distinct bending movement of the body, where the abdomen is bent upwards just after impact (see 6). The presence of bending movements significantly reduced the landing duration and thus could be regarded as a key feature for a fast and secure landing.

Our results also show that bending of the abdomen was also sometimes found in dead and cool insects. This indicates that at least to a certain extend the upwards bending movements is a passive result of the exoskeletons morphology. The reduced occurrence of bending in dead and cool insects could be explained by either different biomechanical properties of the exoskeleton (such as reduced hemolymph pressure, different damping properties of muscles and joints, etc.) or missing or reduced active movements of the exoskeleton supporting the bending. In any case, this biomechanical adaptation helps to reduce rebound and shortens the time to recover. Similar principles can be found in several martial arts, where falling athletes use rolling movements of their body extremities to dissipate energy away from the impact site and reduce impact forces (Groen, Weerdesteyn and Duysens, 2007).

### 3. Conclusion

Our results show that jumping locusts just crash into the substrate and thus rely on their exoskeleton to deal with the impact forces. Using their wings and probably aerodynamic properties of their exoskeleton locusts are able to actively control the impact angle and first point of contact. The adhesive pads and a bending movement of the abdomen help to dissipate energy and ensure a quick recovery after the impact.

## Acknowledgements

The authors would like to acknowledge the help of Florian Hoffmann and René Sonntag in setting up the high-speed-recording equipment. Nikolai Rosenthal and Benjamin Aberle assisted in developing the experimental protocols.

## Author Contributions

SR and JHD conceived and designed the experiments. SR performed the experiments and programmed the analysis sripts. All authors analysed and interpreted the data and wrote the manuscript.

## Data Availability

The data of this study can be made available on individual request.

